# Formation of brain-wide neural geometry during visual item recognition in monkeys

**DOI:** 10.1101/2024.08.05.604527

**Authors:** He Chen, Jun Kunimatsu, Tomomichi Oya, Yuri Imaizumi, Yukiko Hori, Masayuki Matsumoto, Yasuhiro Tsubo, Okihide Hikosaka, Takafumi Minamimoto, Yuji Naya, Hiroshi Yamada

## Abstract

Neural dynamics assumes to reflect computations that relay and transform information in the brain. Previous studies have identified the neural population dynamics in many individual brain regions as a trajectory geometry, preserving a manifest of computational motifs. However, whether these populations share particular geometric patterns across brain-wide neural populations remains unclear. Here, by mapping neural dynamics widely across temporal/frontal/limbic regions in the cortical and subcortical structures of monkeys, we show that 10 neural populations, including 2,500 neurons, propagate visual item information in a stochastic manner. We found that visual inputs predominantly evoked rotational dynamics in the higher-order visual area, TE and its downstream striatum tail, while curvy/straight dynamics appeared frequently downstream in the orbitofrontal/hippocampal network. These geometric changes were not deterministic but rather stochastic according to their respective emergence rates. Our meta-analysis results indicated that visual information propagates as a heterogeneous mixture of stochastic neural population signals in the brain.

## Introduction

Visual inputs activate a large number of neurons in the brain that construct numerous neural networks to process information in an environment (*1–3*). This brain-wide activity change reflects the information processing embedded in each individual neural circuit; however, limitations of spatial and temporal resolution in the measurements of circuitry activity disrupt our understanding of brain-wide visual information processing (*4–9*). Under this limitation, considerable attempts have been made toward understanding how the brain processes information using a variety of developing theoretical frameworks (*10–15*).

One of the analytic frameworks developed within the last decade is state-space analysis (*16*) that provides a mechanistic structure of information processed in the lower-dimensional space of a neural population (*17–19*). This analytical tool identified dynamic neural population structures that reflect information processing for general biological features (*20, 21*) and allowed us to describe those features as a neural geometry in a fine time resolution (*13–15*) in the sub-second order. A large number of unidentifiable neural circuits may process information moment-by-moment (*6*), and they may form a population geometry, such as rotational (*18*), curvy (*19, 22*), or straight geometries (*23*), as the typical and basic features of dynamics. A recent finding suggests that the combination of neural population geometries may be the key to processing information to transform sensory inputs into memory (*24*). Recent studies have extended these analytical frameworks (*25–29*); however, because of poor comparisons among brain-wide neural populations, how the brain processes information in the form of geometry remains elusive.

To examine how brain-wide neural dynamics are formed to process visual information, we accumulated the neural population data of monkeys from four laboratories that contained 10 neural populations, including 2,500 neurons across temporal/frontal/limbic networks (i.e., meta-analysis). We applied targeted dimensional reduction, with a bootstrap resampling technique that detects and replicates neural modulation dynamics in a low-dimensional neural space. Following a parametric bootstrap analysis using the Lissajous curve function, our cross-study comparison revealed that a gradual shift in stochastic neural population signals occurred throughout the temporal-to-frontal brain regions.

## Results

We compared the trajectory geometries across many neural populations widely distributed in the brain from the output brain regions in the ventral visual pathway (*30–33*) and its downstream brain regions that may access memories associated with a visual stimulus. These included ten brain regions that were accumulated from nine monkeys examined in the four laboratories (Table. S1), from the higher-order visual area TE and their downstream brain regions in cortical, subcortical, and limbic structures, such as the temporal/orbitofrontal cortices, striatum, and hippocampus (HPC) (Fig. 1A, No. 1 to 10). A total of 2,500 neurons were accumulated across the four behavioral tasks (Fig. S1), in which visual items provided monkeys with position and/or reward information during the active (Exps. 1 and 3), and passive (Exps. 2 and 4) behavioral responses. Using state-space analysis, we characterized structures of neural population geometry that appeared in the lower dimensional neural space, which describes how neural modulation by the task parameters of interest processes information at the population level (*23, 29*).

**Fig. 1.**
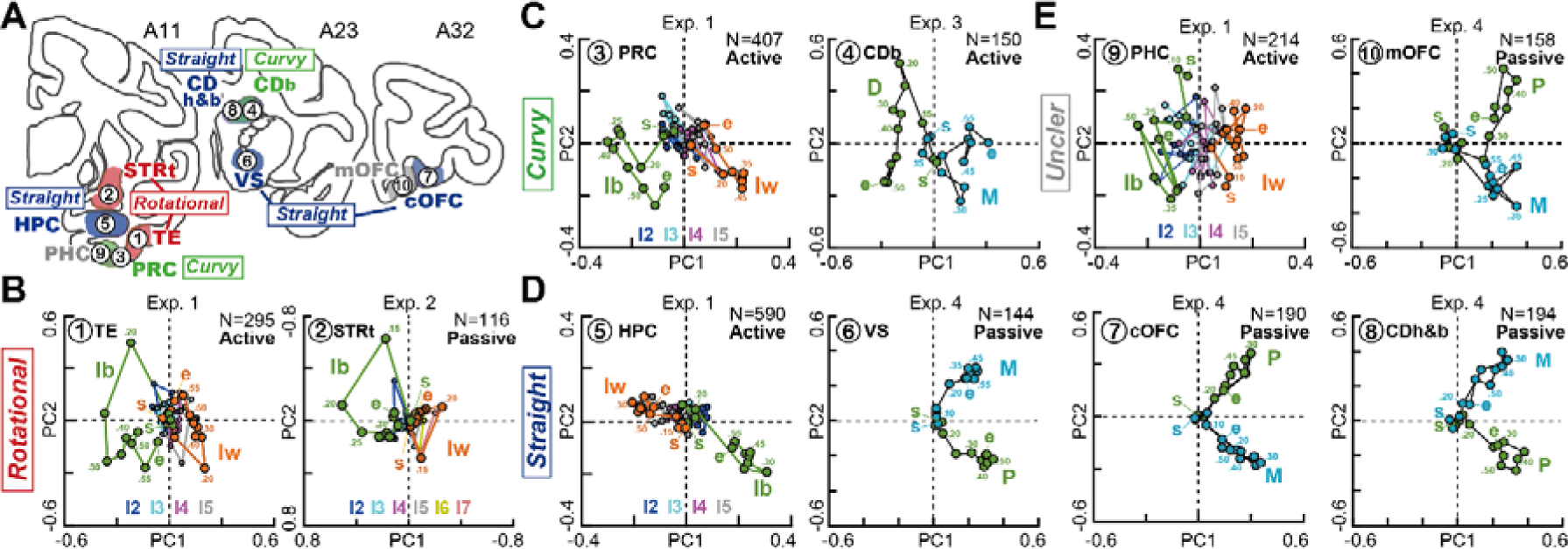
Neural population geometries in the visual memory pathway **A,** Anatomical depiction of neural populations obtained from the 10 brain regions in eight macaques during the four different behavioral tasks in Exps. 1 to 4. **B-E,** Rotational (**B**), curvy (**C**), straight (**D**), and unclear dynamics (**E**) detected by visual inspection. In **A-E**, the 10 brain regions are numbered as follows: 1. TE, 2. STRt, 3. PRC, 4. CDb, 5. HPC, 6. VS, 7. cOFC, 8. CDh&b, 9. PHC, and 10. mOFC. The 0.05 s time bin was used for the dynamics analysis.

We found all three types of geometric patterns, rotational, curvy, and straight geometries, in the top two dimensions (Fig. 1B-D, see Fig. S2 for performance as the percentage of variance explained), including unclear structures based on visual inspection (Fig. 1E). All the 10 neural populations showed a significant structure at the principal component (PC)1-2 plane based on shuffle controls (Fig. S3, *P* < 0.05 for all PC1 and PC2). PC3 did not show statistical significe in some neuronal populations (Fig. S3, Exps. 2 and 4; compare the black and gray dots for each PC). These identified geometric structures appeared to be distributed from complex to simple, reflecting the circuit distance to the visual input (Fig. 1). For example, TE (Fig. 1B, No. 1, A11 plane in A) and its downstream region striatum tail (STRt) (Fig. 1B, No. 2, A11 plane in A), showed rotational geometries during visual item recognition. In more detail, at the beginning of information processing after the visual item presentation (Fig. 1B left, see green **s** at time = 0), the neural state was positioned at around the center of the PC1-2 plane, and then rotated at approximately 0.2 s and started to move the second quadrant with the counterclockwise rotation going back close to the initial point (see green **e** 0.6 s after visual item onset). An opposite rotation was observed for the worst visual item, with a smaller change (orange). These rotational structures were also observed at the STRt on a similar timescale (Fig. 1B, right).

In the downstream brain regions, such as the perirhinal cortex (PRC) and caudate body (CDb), rotational or curvy dynamics were observed (Fig. 1C, No. 3, PRC, A11 plane in A and No. 4, CDb, A23 plane in A), which were characterized by a half rotation ending at the opposite neural space and end points deviating from the initial point, although it was unclear whether they showed rotational or curvy dynamics. In contrast, straight dynamics were observed in brain regions far away from the visual inputs in the HPC (Fig. 1A, No. 5 in the A11 plane, Fig. 1D) and central part of the orbitofrontal cortex (cOFC) (Fig. 1A, No. 7 in the A32 plane, Fig. 1D). In addition, the ventral striatum (VS) showed straight dynamics (Fig. 1A, No. 6 in the A23 plane, Fig. 1D), although some structures could not be clearly determined (Fig. 1E, parahippocampal cortex, PHC, medial orbitofrontal cortex, mOFC). The straight dynamics also showed a geometric change back and forth along the straight trajectory (Fig. 1D). These qualitative observations based on visual inspection suggest that neural population structures may change dynamically through visual recognition process, and the shift of neural population geometries might occur throughout the cortical and sub-cortical structures across the temporal/frontal/limbic network.

### Evaluation of geometric patterns based on their selected features

To quantify these geometric patterns occurring at approximately half a second, we estimated indices for the characteristics of dynamic neural changes in the low-dimensional neural state (Fig. 2A). They were as follows: accumulated angle difference weighted by deviance, Σ*d θ*, which reflects a measure similar to the centrifugal force (see Materials and Methods for details, Fig. 2A, top); mean distance of vectors (*d*, Fig. 2A, top); rotational speed (*Θ*/0.1 s, Fig. 2A, bottom); and distance between start and end of trajectory (*d*_S-E_, Fig. 2A, bottom). Following the replication of the neural population geometries based on the bootstrap resampling technique (see Materials and Methods), we calculated these parameter values for each replicated neural population.

**Fig. 2.**
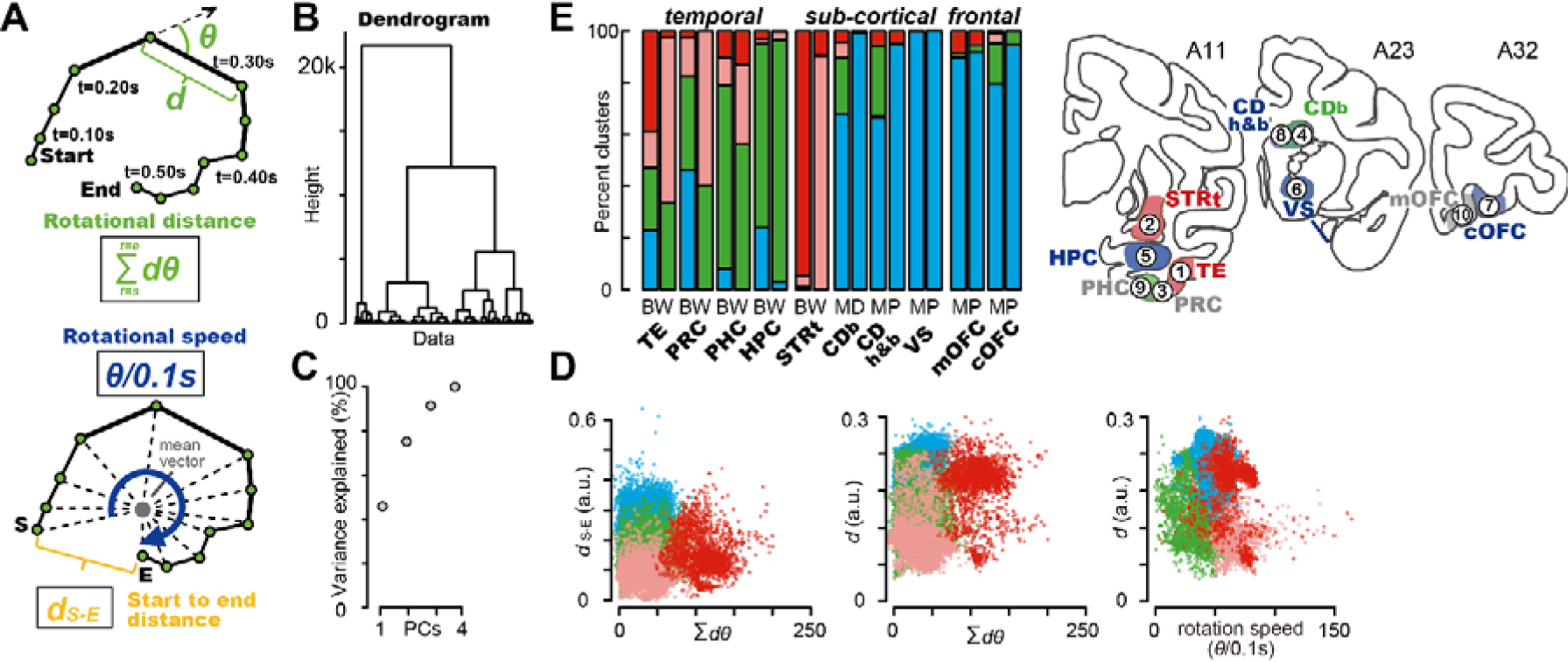
Quantitative evaluation of geometric structures according to the rotational features. **A,** Schematic depictions of the estimation of accumulated angle difference weighted by the deviance, Σ*d θ*. The accumulated angle difference indicates the degree of geometric change in terms of the rotational force across time. Vector distance (*d*), rotational speed (*θ*/0.1s), and start to endpoint distance (*d*_S-E_) were also estimated. **B,** Dendrogram estimated from these four parameter values based on bootstrap resampling across 10 neural populations. **C,** Percentage of variance explained by PCA of bootstrap resampling data across 10 neural populations. **D,** Clusters detected among the four parameters based on the PCA. **E,** Percentage of the identified clusters in each of the 10 brain regions. Each neural population contained two components of neural information: the best and worst conditions (BW) in Exps. 1 and 3, magnitude and delay of the rewards (MD) in Exp. 2, and magnitude and probability of rewards (MP) in Exp. 4.

We found that across the 10 neural populations, these indices captured geometric features to some extent, similar to the rotational geometries observed in the TE and STRt (Fig. 2B-E). For instance, identified clusters based on the dendrogram and principal component analysis (PCA) (Fig. 2B-C) showed that they possess a rotational characteristic with high rotational speed (Fig. 2D, right, red), large Σ*d θ* (Fig. 2D, middle, red), large *d* (Fig. 2D, middle, red), and small *d*_S-E_ (Fig. 2D, left, red), which occupy more than 90% of the STRt population in the best item condition (Fig. 2E, see also Fig. 1B, right green trajectory). A smaller rotational structure characterized by smaller values of Σ*d θ* was also captured by another cluster (Fig. 2D and E, shallow red), which occupied approximately 90% of the STRt population in the worst item condition (Fig. 2E, see also Fig. 1B right, orange trajectory). These rotational features were observed in other temporal brain regions (Fig. 2E, see reddish, more than 50% in TE, 20-40% in PRC and PHC), but merely observed at the frontal/limbic brain regions, such as HPC (less than 10% in all remaining brain regions). In contrast, curvy/straight dynamics were observed in other clusters in the downstream brain regions (Fig. 2E, green and blue).

Collectively, in each of the 10 brain regions, a cluster with high rotational speed occupied the STRt and half of the TE populations (Fig. 2D, reddish), while the curvy/straight dynamics occupied most of the replicates in the remaining cortical and subcortical regions (Fig. 2D and E, blue and green).

### Parameterization of geometric patterns using Lissajous curve function

To parameterize these geometric features in more detail, we fitted the Lissajous curve function (*34*) to the replicated data, which can mathematically capture all rotational, curvy, and straight dynamics by this single functinon. In the Lissajous function, any two-dimensional geometric features represented by *F(x, y)* are captured using the following equations:

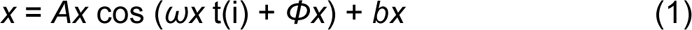

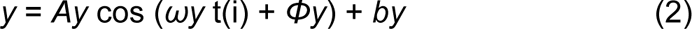

where *ω* and *Φ* represent cycle of the rotation and their deviance as a function of time, t(i). t(i) takes the values from 0 to 0.6 s in all the four experiments, and thus, one cycle of the trajectory is represented as 0 to 3.33 π for *ω*. For the horizontal and vertical axes, *Ax* and *Ay* define the amplitude of the trajectory, respectively, whereas *bx* and *by* determine its location. In this function, rotational dynamics is represented by the same *ω* among *x* and *y* formulas, and 0.5 π cycle differences in *Φ* values between *x* and *y* formulas (Fig. 3A, left, see also Fig. S4A for more details). We note that this rotational example represents less than one cycle due to *ωx* = 3.0 π. In contrast, straight dynamics are represented with the same *ω* and also same *Φ* values between the two formulas (Fig. 3A, right, see also Fig. S4C). Curvy dynamics are represented with some difference of *ω* and same *Φ* values (Fig. 3A, middle, see also Fig. S4B). We fitted this Lissajous curve function to each of the 20,000 bootstrap replicates derived from the 10 neural populations (see Materials and Methods, 1000 replicates times 10 populations times two conditions). For instance, three replicated examples obtained from the HPC population were well captured by the Lissagous curve function, as rotational-to-straight trajectories (Fig. 3B). We obtained all the estimates for these parameters (Fig. 3C), and thereafter, applied clustering to these data (Fig. 3D-F) to identify geometry types as a function of the Lissajous curve parameters.

**Fig. 3.**
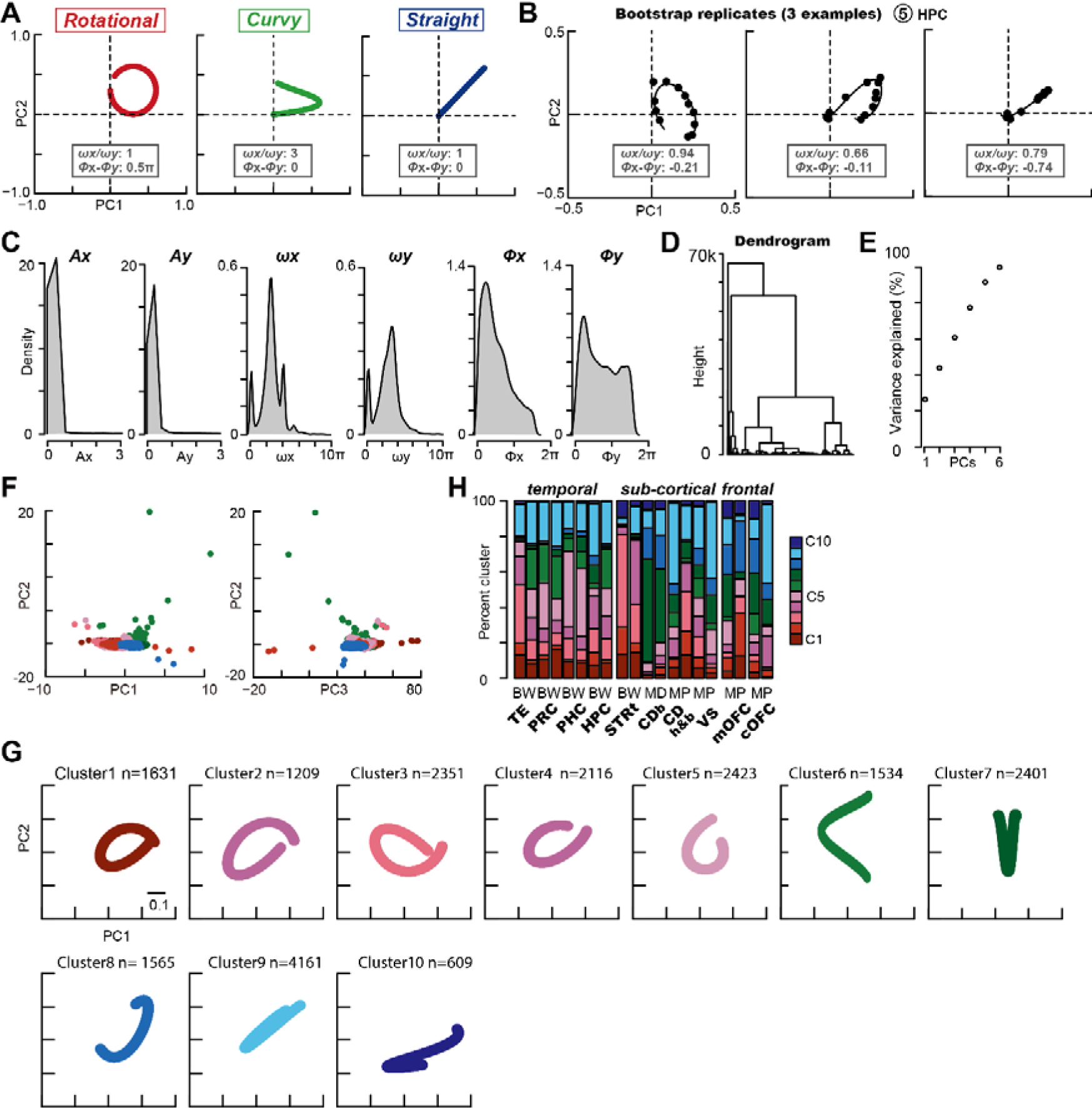
Quantitative evaluation of geometric structures using the Lissajous curve function. **A,** Schematic depictions of trajectory geometries using Lissajous function parameters. For all figures, *ωx* and *ωy* are 3 π. **B,** Three examples of bootstrap replicates for the HPC population fitted by the Lissajous function. *L* indicates the maximum loglikelihood. Estimated parameters were as follows: left, *ωx*, 2.78 π, *Φx*, −0.11, *L*, 34.6, *ωy*, 2.96 π, *Φy*, 0.10, *L*, 31.6; middle, *ωx*, 2.51 π, *Φx*, −0.03, *L*, 24.2, *ωy*, 3.82 π, *Φy*, −0.08, *L*, 32.7; right, *ωx*, 2.21 π, *Φx*, −0.16, *L*, 26.9, *ωy*, 2.78 π, *Φy*, 0.58, *L*, 29.1. **C,** Probability density estimated for Lissajous parameters obtained from bootstrap replicates across 10 neural populations times two conditions. **D,** Dendrogram estimated from Lissajous parameter values based on bootstrap resampling across 10 neural populations times two conditions. **E,** Percentage of variance explained by PCA of bootstrap resampling data across 10 neural populations times two conditions. **F,** Clusters determined using PCA. Data are shown for PC1 to 3. **G,** Reconstructed trajectory in each cluster based on bootstrap resampling. The trajectories in clusters 1–10 were drawn using the median values of the Lissajous parameters in each cluster. **H,** Percentage of clusters in each of the 10 brain regions times two conditions. BW: best and worst conditions. MD: magnitude and delay conditions. MP: magnitude and probability conditions.

We found that the rotational dynamics (Fig. 3G-H, clusters C1-C5, reddish) appeared at the TE and STRt, which occupied high percentages of these neural populations (Fig. 3H, approximately 70%), and they were also observed in more than 50% of temporal brain regions. Cluster 5 seemed to have the intermediate characteristics between rotational and curvy structures; if we define this cluster 5 as the curvy one, the rotational percentage becomes low in the PRC, PHC, and HPC (30-40%), but not in the STRt (50-70%). Curvy structures were predominantly observed in the CDb population in more than 40% (Fig. 3G and H, Clusters 6 and 7, greenish). These clusters were note observed deterministically but rather stochastically, as also seen in the predominant percentages of intermediate features between curvy and straight dynamics (Fig. 3G and H, Cluster 8). Straight dynamics were predominant at the frontal brain regions, while they were also observed at the temporal cortices (Fig. 3G and H, Clusters 8-10, bluish). Even with the STRt, the neural population contained curvy or straight dynamics in more than 20%. These heterogeneous mixtures of replicated signals in each population suggested that neural dynamics emerged in a stochastic manner with a functional gradient in the temporal/frontal/limbic networks of cortical and subcortical structures. The brain-wide neural population may propagate item information as a heterogeneous mixture at approximately half a second.

## Discussion

Collectively, our results revealed that parts of the orbitofrontal cortex (cOFC and mOFC) and their target subcortical brain region VS predominantly showed curvy, straight, and intermediate dynamics (Fig. 3G-H). These dynamics exhibited maximum modulation at approximately 0.3 sec after visual onset, except in the slowest VS dynamics (Fig. 1D). Rotational dynamics were observed at the temporal cortices and their connected striatal regions, at the relatively shorter latency around 0.2 s when rotation started (Fig. 1B). Remarkably, the rotational dynamics were observed at different neural proportions of replicated population in a stochastic manner (Fig. 3H), whereas different monkeys performed active (Exps. 1 and 3), and passive (Exps. 2 and 4) behavioral tasks. In contrast, straight dynamics started its geometric change approximately after 0.2 s of the visual onset (Fig. 1D, see the distance between initial point S and 0.2 s location), indicating that they follow the rotational dynamics. Taken together, these three dynamics were distinctive in terms of geometric patterns and their dynamic changes over time (Fig. 4B for summary), in which a rotational/curvy change was followed by a change in straight dynamics.

**Fig. 4.**
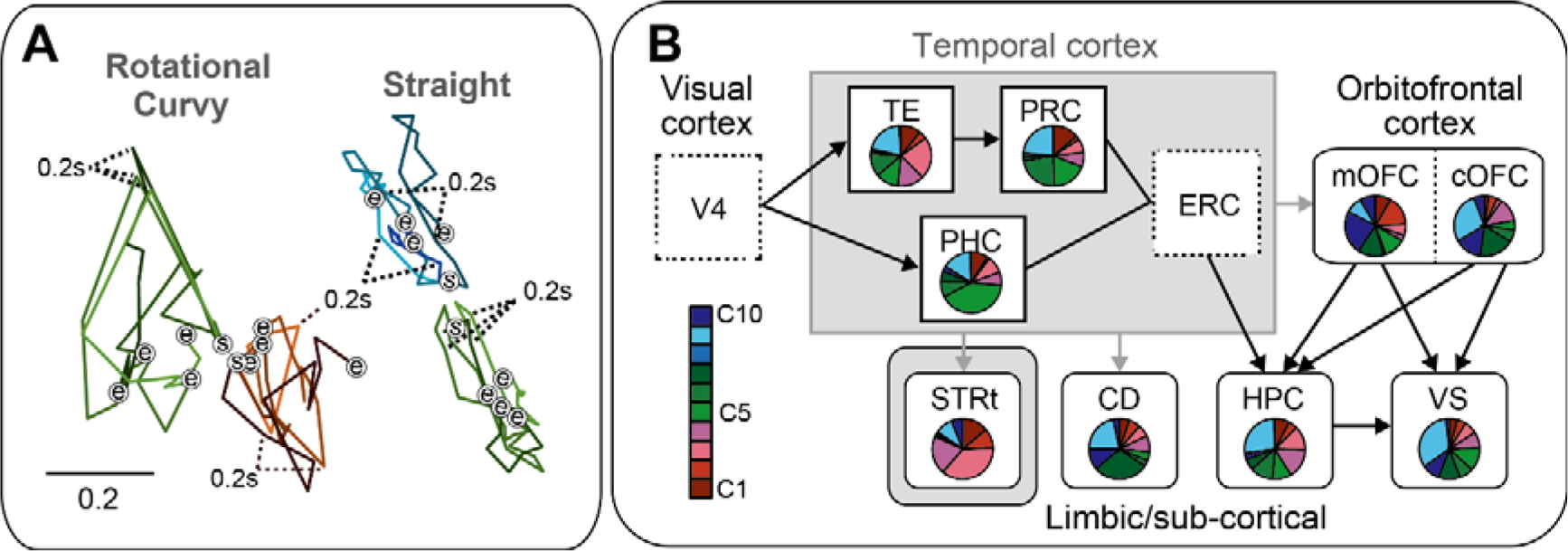
Summary of the observed dynamics and anatomical connections in the visual memory pathway. **A,** Geometries depicted in the same arbitrary scales on the PC1-2 plane for the eight neural populations shown in Fig. 1B-D. The start of the trajectory (S) is aligned to describe each trajectory. e indicates the end of the trajectory at 0.6 s. **B,** Proportion of the clusters defined in each of the 10 brain regions are described with the anatomical connection. Reddish: rotational, greenish: curvy, bluish: straight dynamics. Data from CDh&b and CDb are merged (CD).

A previous study showed that rotational dynamics have been uncovered broadly in the primary sensory (*24, 35*) and motor (*36*) cortices, which are closer to the inputs and outputs of the brain, such as motor unit activity (*37*). Other studies have shown that the prefrontal cortex (*38, 39*) and parietal cortex (*22, 35*) exhibit curvy dynamics. We found curvy dynamics in the CDb, where action information was transferred from the cortices (*40*). These characteristic differences in the individual previous studies were not deterministic, similar to the stochastic differences among the populations observed in the present study, which may reflect the neural computations processed in the brain.

In the present study, we specifically focused on neural dynamics in two core areas: i) low-dimensional geometries and ii) neural modulation dynamics. First, although these data were obtained from four different laboratories using distinctive behavioral tasks during the passive and active responses of monkeys, the low-dimensional features of neural dynamics are thought to be preserved across mammals in the brain-wide network (*41, 42*). A recent study provides clear evidence that different animal species share and preserve their neural geometries during behavioral tasks (*41*). It is suggested that a low-dimensional manifold in the neural state space might be one of the representational states of biologically relevant information, similar to many combinations of physical properties in the world (*43*). Second, the dynamics in neural modulations examined here are comparable with those using the standard analytic frameworks in the rate coding model, which has provided a huge amount of knowledge corresponding to low-dimensional neural activity modulation in the literature, such as the Gabor function in the visual cortices (*44, 45*), movement direction and muscle force in the motor cortex (*46, 47*), reward value in the parietal cortex (*48*), and comprehension of the location of animals during navigation in the HPC (*49*). Thus, in our analysis, the dynamics of these well-known brain region features were compared as the geometric patterns across neural populations during visual recognition (*29*).

One concern with our approach is that there may be limitations in data interpretation in terms of data sharing and comparisons across different behavioral tasks and different individual animals. Is it possible to compare the neural population trajectory using accumulated data across animals and tasks with a certain analytical tool? In previous literature, when we analyzed neural modulation using a linear regression model, comparisons of firing rate modulations were accepted by most neurophysiologists for different animals as well as different behavioral tasks. In one study, a challenge was made to compare the neural modulation dynamics as the trajectory geometry between different laboratories’ data (*50*). Thus, the neural trajectory would be comparable among our shared data with greater deliberation.

Our findings would add to the emerging literature describing how visual inputs alter the brain-wide neural dynamics associated with visual memories by connecting neural geometry types and their alignments across many brain regions. Previous studies have shown the existence of different types of neural population dynamics in each individual study (*22, 24, 35, 36, 38, 39*). Although some of these dynamics may reflect task demand, as observed in the dorsolateral prefrontal cortex (*38, 51*), they are difficult to disentangle from changes in behavior and neural activity levels and may involve some transformation of information for behavioral responses (*52*). It is possible that the dorsal and motor-related brain regions have this type of flexibility in their dynamics, as partly observed in this study in the CDb, where curvy dynamics were predominantly observed (Fig. 4B).

Our results raise the possibility that geometric features determine the important neural mechanisms widely observed in the brain. For instance, the stochastic gradient relative distance to the visual input may reflect the dynamics of the neural circuitry in half a second (Fig. 4B). The unidimensional straight dynamics in the hippocampal-frontal circuitry may reflect memory access during visual recognition, such as location and reward. The rotational dynamics might reflect the visual recognition process, during which recurrent feedback signals change the circuit dynamics. Future studies should test the underlying mechanisms and define whether engagement is best considered a change in behavior and/or task context for whole-brain neural population activity. Regardless of the mechanism, the shift of modulation structures in the lower-dimensional neural space could play a fundamental role in brain-wide information processing, such as transforming visual feature recognition to memory access.

## Materials and Methods

### Subjects and experimental procedures

Nine rhesus monkeys were used in the present study (Exp. 1: *Macaca mulatta,* A, 9.3 kg, male; *Macaca mulatta,* D, 9.5 kg, male; Exp 2: Macaca mulatta, WK, 12.0 kg, male; Macaca mulatta, SP, 7.0 kg, male; Exp 3: *Macaca mulatta,* BI, 8.2 kg, male; *Macaca mulatta,* FG, 11.0 kg, male; *Macaca mulatta,* ST, 5.2 kg, male; Exp 4: *Macaca mulatta,* SUN, 7.1 kg, male; *Macaca fuscata*, FU, 6.7 kg, female). All experimental procedures were approved by the Institutional Animal Care and Use Committee of Laboratory Animals approved by Peking University (Exp. 1, project number Psych-YujiNaya-1), and Animal Care and Use Committee of the National Eye Institute, and complied with the Public Health Service Policy on the Humane Care and Use of Laboratory Animals (Exp. 2, protocol number NEI-622), the Animal Ethics Committee of the National Institutes for Quantum Science and Technology (Exp. 3, protocol no. 11-1038-11), and the Animal Care and Use Committee of the University of Tsukuba (Exp. 4, protocol no H30.336). All procedures were performed in compliance with the US Public Health Service Guide for the Care and Use of Laboratory Animals.

### Behavioral task

#### Exp. 1. Item-location-retention (ILR) task

The animals performed the task under dim light conditions in an electromagnetically shielded room. The task started with an encoding phase, which was initiated by the animal pulling a lever and fixating on a white square (0.6°) presented within one of four quadrants at 12.5° (monkey A) or 10° (monkey D) from the center of the touchscreen (3M^TM^ MicroTouch^TM^ Display M1700SS, 17 in), situated approximately 28 cm from the subjects. The eye position was monitored using an infrared digital camera with a sampling frequency of 120 Hz (ETL-200, ISCAN). After fixation for 0.6 s, one of six items (3.0° for monkey A and 2.5° for monkey D, radius) was presented in the same quadrant as a sample stimulus for 0.3 s, followed by another 0.7 s fixation on the white square. If the fixation was successfully maintained (typically < 2.5°), the encoding phase ended with the presentation of a single drop of water.

The encoding phase was followed by a blank interphase delay interval of 0.7–1.4 s during which no fixation was required. The response phase was initiated using a fixation dot presented at the center of the screen. One of the six items was then presented at the center for 0.3 s, as a cue stimulus. After another 0.5 s delay period, five disks were presented as choices, including a blue disk in each quadrant and a green disk in the center. When the cue stimulus was the same as the sample stimulus, the animal was required to make a choice by touching the blue disk in the same quadrant as the sample (i.e., the match condition). Otherwise, the subject was required to choose the green disk (i.e., non-match condition). If the animal made the correct choice, four–eight drops of water were provided as a reward; otherwise, an additional 4 s was added to the standard inter-trial interval (1.5–3 s). The number of reward drops was increased to encourage the animal to maintain good performance in the latter phase of a daily recording session, which was typically conducted in blocks (e.g., a minimal set of 60 trials with equal numbers of visual items presented in a match/nonmatch condition). During the trial, a large gray square (48° on each side) was presented at the center of the display as a background. At the end of the trial, all stimuli disappeared, and the entire screen displayed a light red color during the inter-trial interval. The start of a new trial was indicated by the reappearance of a large gray square on the display, at which point the monkey could pull the lever, triggering the appearance of a white fixation dot.

In the match condition, sample stimuli were chosen pseudo-randomly from six well-learned visual items, and each item was presented pseudo-randomly within four quadrants, resulting in 24 (6 × 4) configuration patterns. In the non-match condition, the location of the sample stimulus was randomly chosen from the four quadrants, and the cue stimulus was randomly chosen from the remaining five items that differed from the sample. The match and non-match conditions were randomly presented in a ratio of 4:1, resulting in 30 (24 + 6) configuration patterns. The same six stimuli were used during all the recording sessions.

#### Exp. 2. Scene-based object-value task

Animals learned the scene-object associations. After the monkeys fixated on the red-square fixation point on the scene image for 0.6–1 s, the fixation cue disappeared, and two visual items (objects of different values) appeared simultaneously in a different hemifield (for training and neuronal testing) or the same hemifield (for pharmacological experiments). A reward was given after the monkeys made a saccade to the stimulus and maintained fixation for 0.2 s. Half of the fractal visual items were associated with a large reward (0.3 mL), and the other half were associated with a small reward (0.1 mL). This reward association changed depending on the scene (Fig. S1 D). *Passive Viewing Task.* One of the two scene images was presented for 0.8 s randomly. If the monkey fixated on a central red square, two to four fractals were presented sequentially on the scene image within the neuron’s receptive field (presentation time, 0.4 s; interstimulus interval, 0.4 s; Fig. S1C). A liquid reward (0.2 mL) was delivered 0.3 s after the last object was presented. Thus, reward occurrence was not associated with any of the visual items. Each item was presented at least seven times per session.

#### Exp. 3. Delayed reward tasks

The monkeys were seated on a primate chair inside a dark, sound-attenuated, electrically shielded room. A touch-sensitive bar was mounted on the chair. The visual stimuli were displayed on a computer video monitor placed in front of the animals. Each of the six cues was associated with a combination of reward size (1 drop; 3 or 4 drops) and reward delay (0, 3.3, and 6.9 s). The trials began when the monkey touched the bar. A visual cue appeared, and the monkey released a bar when a red spot (waiting signal) turned green (go signal) after a variable interval. If the monkey released the bar 0.2–1 s after this go signal, the trial was considered correct and the spot turned blue (correct signal). A liquid reward of a small (1 drop, approximately 0.1 mL) or large amount (3 drops, except for monkey BI, 4 drops) was delivered immediately (0.3 ± 0.1 s) or with an additional delay of either 3.3 ± 0.6 s or 6.9 ± 1.2 s after correct release of the bar. The cues were chosen with equal probability and were independent of the preceding reward condition. Anticipatory bar releases (before or no later than 0.2 s after the appearance of the go signal) and failure to release the bar within 1 s of the appearance of the go signal were counted as errors. In the error trials, the trial was terminated immediately, all visual stimuli disappeared, and following inter trial interval (1 s), the trial was repeated; that is, the reward size/delay combination remained the same as that in the error trial. Behavioral control and data acquisition were performed using a real-time experimentation system (REX) (*53*). The Neurobehavioral Systems Presentation software was used to display the visual stimuli (Neurobehavioral Systems).

#### Exp. 4. Cued lottery tasks

The animals performed one of two visually cued lottery tasks: a single-cue or a choice task. Neuronal activity was only recorded during the single-cue task.

Animals performed the task under dim lighting conditions in an electromagnetically shielded room. Eye movements were measured using a video camera system at 120 Hz (EyeLink, SR Research). Visual stimuli were generated using a liquid-crystal display at 60 Hz, placed 38 cm from the monkey’s face when seated. At the beginning of the single-cue task trials, the monkeys had 2 s to align their gaze within 3^°^ of a 1^°^-diameter gray central fixation target. After a fixation for 1 s, a pie chart was presented for 2.5 s, to provide information regarding the probability and magnitude of rewards in the same location as the central fixation target. The probability and magnitude of the rewards were associated with the number of blue and green 8^°^ pie chart segments, ranging from 0.1 to 1.0 mL in 0.1 mL increments for magnitude, and 0.1 to 1.0 in 0.1 increments for probability. Following a 0.2 s interval from the removal of the pie chart, a 1 kHz or 0.1 kHz tone of 0.15 s duration was provided to indicate reward or no-reward outcomes, respectively. After a 0.2 s interval following the high tone, a fluid reward was delivered, whereas no rewards were delivered following the low tone. An inter-trial interval of 4–6 s was used. During the choice task, animals were instructed to choose one of two peripheral pie charts, each of which indicated either the probability or magnitude of an upcoming reward. The two target options were presented for 2.5 s at 8^°^ to the left or right of the central fixation location. The animals received a fluid reward as indicated by the green pie chart of the chosen target, with the probability indicated by the blue pie chart. Otherwise, no reward was delivered.

A total of 100 pie charts composed of 10 levels of probability and magnitude of rewards were used in the experiments. In the single-cue task, 100 pie charts were presented once in random order. In the choice task, two pie charts were randomly assigned to the two options. During one electrophysiological recording session, approximately 30–60 trial blocks of the choice task were interleaved with 100–120 trial blocks of the single-cue task.

### Electrophysiological recordings and data preprocessing

#### Exp. 1

To record the single-unit activity, we used a 16-channel vector array microprobe (V1 × 16-Edge, NeuroNexus), 16-channel U-Probe (Plexon), tungsten tetrode probe (Thomas RECORDING), or single-wire tungsten microelectrode (Alpha Omega).

Electrophysiological signals were amplified, bandpass-filtered (200–6000 Hz), and monitored. Single-neuron activity was isolated based on spike waveforms, either online or offline. For both clustering and offline sorting, the activities of all single neurons were sampled when the activity of an isolated neuron demonstrated a good signal-to-noise ratio (>2.5). The signal-to-noise ratio was visually checked by calculating the range of background noise against the spike amplitude, which was monitored online using the OmniPlex Neural Data Acquisition System, or offline using the sorter software Plexon. The recorded neurons were not blinded. The sample sizes required to detect the effect sizes (numbers of recorded neurons, recorded trials in a single neuron, and monkeys) were estimated based on previous studies (*31, 54*). Neural activity was recorded during 60–240 trials of the ILR task. We recorded 590 hippocampal neurons, among which the recording sites appeared to cover all subdivisions (i.e., the dentate gyrus, CA3, CA1, and subicular complex).

#### Exp. 2

We used conventional techniques to record the single-neuron activity in the STRt, including the caudate and putamen tails. A tungsten microelectrode (1–3 MΩ Frederic hair; 0.5-1.5 MΩ Alpha Omega Engineering) was used to record single-neuron activity. The recording site was determined using a grid system that allowed electrode penetration at 1 mm intervals. We amplified and filtered (0.3 to 10 kHz; Model 1800, A-M Systems; Model MDA-4I, BAK) signals obtained from the electrodes and collected at 1 kHz. Single neurons were isolated online using custom voltage–time window discriminator software (Blip; available at http://www.robilis.com/blip/). The presumed medium spiny neurons were identified based on their low baseline activity (<3 spikes/s) and broad action potentials (*55*). The recorded neurons were not blinded. The sample sizes required to detect the effect sizes (numbers of recorded neurons, recorded trials in a single neuron, and monkeys) were estimated based on previous studies (*56, 57*). Neural activity was recorded during 10–30 trials of the *passive viewing task*. We recorded 115 medium spiny neurons in the STRt. In Exp. 2, only a single-neuron recording was performed online. We note that we termed the scene and object for two visual stimuli in our previous study (*58*), but here we termed them scene and item.

#### Exp. 3

Conventional techniques were used to record single-neuron activity in the dorsal part of the head of the caudate nucleus (CD). A tungsten microelectrode (1.1–1.5 MΩ, Microprobes for Life Science; 1.0 MΩ, Alpha Omega Engineering Ltd.) was used to record single-neuron activity. The electrophysiological signals were amplified and monitored using a TDT recording system (RZ2, Tucker-Davis Technologies, TDT). Single-neuron activity was manually isolated based on the online spike waveforms. The activity of all single neurons was sampled from the activity of presumed projection neurons, which are characterized as having a low spontaneous discharge rate (<2 spikes/s) outside the task context and exhibiting phasic discharges in relation to one or more behavioral task events (Yamada et al, 2016). Neural activity was recorded during 100–120 trials per block in the delayed-reward task. We recorded the CD of the left or right hemisphere in each of the three monkeys in the experiment, with 150 CD neurons (51, 31, and 68 from the BI, FG, and ST, respectively).

#### Exp. 4

Conventional techniques were used to record single-neuron activity in the DS, VS, cOFC (area 13M), and mOFC (area 14o). A tungsten microelectrode (1–3 MΩ, FHC) was used to record single-neuron activity. Electrophysiological signals were amplified, band-pass filtered (50–3,000 Hz), and monitored using a TDT recording system (RZ5D, Tucker-Davis Technologies, TDT). Single-neuron activity was manually isolated based on the online spike waveforms. The activity of all single neurons was sampled when the activity of an isolated neuron demonstrated a good signal-to-noise ratio (>2.5). The signal-to-noise ratio was calculated online as the ratio of the spike amplitude to the baseline voltage range on the oscilloscope. The recorded neurons were not blinded. The sample sizes required to detect the effect sizes (numbers of recorded neurons, recorded trials in a single neuron, and monkeys) were estimated based on previous studies (*59–61*). Neural activity was recorded during 100–120 trials of the single-cue task. Neural activity was not recorded during selection trials. We recorded the neurons of a single right hemisphere in each of the two monkeys: 194 DS neurons (98 and 96 from monkeys SUN and FU, respectively), 144 VS neurons (89 SUN and 55 FU), 190 cOFC neurons (98 SUN and 92 FU), and 158 mOFC neurons (64 SUN and 94 FU). In Exp. 1, only a single-neuron recording was performed online.

### Statistical analysis

For statistical analysis, we used the statistical software package MATLAB (MathWorks, Exps. 1 and 2), and R (Exps. 3 and 4) for conventional analyses such as linear regression and ANOVA. To analyze the regression matrix using PCA, we used R software. All statistical tests for the neural analyses were two-tailed.

### Behavioral analysis

No new behavioral results were included; however, the procedure for the behavioral analysis was as follows:

#### Exp. 1

We previously reported that two monkeys learned to retain the item and location information of a sample stimulus (*62*). Here, we describe the analysis steps used to check whether the monkey used both item and location information to perform the task.

To examine this, we compared the animals’ actual correct rates during the recording to random correct rates (chi-square test). The ILR response phase had five options, resulting in a 20% random correct rate. If the animal used an incorrect strategy, such as only retaining the location information of the sample stimulus and ignoring the item information, the correct rate for the match condition would be 100% and that for the nonmatch condition would be 0. Based on the above considerations, we examined the correct rates of the two animals in the match and nonmatch conditions, respectively. In general, the average correct rates for both animals in the match and nonmatch conditions were well above chance levels after training.

#### Exp. 2

We previously reported that two monkeys switched their behavior depending on the value of the item based on the scene (*58*). Here, we describe how to check whether the monkey learned both the scene and item information. We calculated the correct rate for the scene-based object-value task. Because the two scenes appear in random sequences, the monkey must switch object choice if the scene has changed. After performing more than 160 trials, the correct rate reached a plateau above chance. The monkey was able to switch object choices immediately after the scene changed. Once the monkeys learned this extensively, their choice behavior became automatic, as the choice tended to occur even when the reward was not delivered after saccades to high-valued items according to the scene.

#### Exp. 3

We previously reported that the three monkeys behaved based on temporally discounted values that integrated both delay and reward size information provided by visual stimuli (*63*). Here, we describe an analysis to check how monkeys discount reward values for delay and reward information. Error rates in task performance were calculated by dividing the total number of errors by the total number of trials for each reward condition and then averaged across all sessions. The average error rates were fitted to the inverse function of reward size with hyperbolic temporal discounting: *E* = 1+*kD*/ *aR* (*E*: average error rates, *D*: delay, *R*: reward size, *k*: discounting factor, *a*: incentive impact), and exponential temporal discounting: *E* = *e*^-*kd*^/*aR*. We used the ‘optim’ function in R, evaluated the goodness of fit of the two models by least-squares minimization, and compared the models by leave-one-out cross-validation as described previously (Minamimoto et al., 2009).

#### Exp. 4

We previously reported that monkey behavior depends on expected values, defined as the probability time magnitude (*23*). We described the analysis steps to check whether the monkey’s behavior reflected task parameters, such as reward probability and magnitude. Importantly, we showed that the monkey’s choice behavior reflected the expected values of the rewards, that is, the probability multiplied by the magnitude. For this purpose, the percentage choosing the right-side option was analyzed in the pooled data using a general linear model with a binomial distribution:

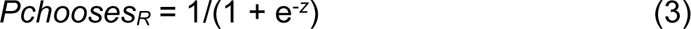

where the relationship between *Pchooses_R_* and *Z* is given by the logistic function in each of the following three models: number of pie segments (M1), probability and magnitude (M2), and expected values (M3).

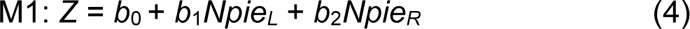

where *b*_0_ is the intercept, and *Npie_L_* and *Npie_R_* are the number of pie segments contained in the left and right pie chart stimuli, respectively. The values of *b*_0_ to *b*_2_ are free parameters and estimated by maximizing the log likelihood.

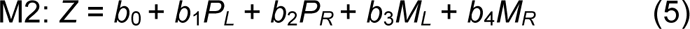

where *b*_0_ is the intercept; *P_L_* and *P_R_* are the probabilities of rewards for the left and right pie chart stimuli, respectively; and *M_L_* and *M_R_* are the magnitudes of rewards for the left and right pie chart stimuli, respectively. The values of *b*_0_ to *b*_4_ are free parameters and estimated by maximizing the log likelihood.

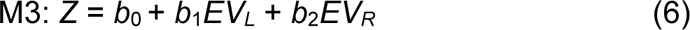

where *b*_0_ is the intercept and *EV_L_* and *EV_R_* are the expected values of rewards as probability multiplied by magnitude for the left and right pie chart stimuli, respectively. The values of *b*_0_ to *b*_2_ are free parameters and estimated by maximizing the log likelihood. We identified the best model to describe monkey behavior by comparing goodness-of-fit based on Akaike’s information criterion and Bayesian information criterion (*64*).

### Neural analysis

Peri-stimulus time histograms were constructed for each single-neuron activity aligned at the onset of the visual stimulus. Average activity curves were smoothed for visual inspection using a Gaussian kernel (σ = 20, 15, 10, and 50 ms in Exps. 1–4, respectively), whereas the Gaussian kernel was not used for statistical tests.

To ensure that the four different datasets were as fair as possible, we used the same criteria to analyze the neural activity. For the neural analyses, we used the following four criteria: 1) the same analysis window size, 2) visual response within a short time (0.6 s), 3) neural modulations detected at the same significance level (P < 0.05), and 4) a general linear model (ANOVA in Exps. 1 and 2 and the linear regression in Exp. 3 and 4). The details of these analytical procedures for the rate coding and dynamic models are shown below.

### Rate-coding model: Conventional analyses to detect neural modulations in each neuron

#### Exp. 1

For neural responses during the encoding phase after the sample presentation, we evaluated the effects of “item” and “location” for each neuron using two-way ANOVA (*P* < 0.05 for each). We analyzed neurons that were tested in at least 60 trials (10 trials for each stimulus and 15 trials for each location). On average, we tested 100 trials for each neuron. These results have been previously reported (*62*).

#### Exp. 2

For neural responses during the appearance of the visual item, we evaluated the effects of “item” and “scene” for each neuron using paired t-test (*P* < 0.05 with Bonferroni correction). These results have been previously reported (*58*).

#### Exp. 3

The neural discharge rates (*F*) were fitted using a linear combination of the following variables:

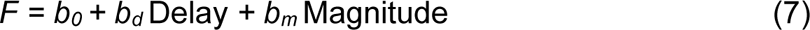

where Delay and Magnitude are the delay and magnitude of the reward, respectively, as indicated by the visual stimulus. *b*_0_ is the intercept. If *b_d_* and *b_m_* were not zero at *P* < 0.05, the discharge rates were regarded as being significantly modulated by that variable. These results have been previously reported (*63*).

#### Exp. 4

The neural discharge rates (*F*) were fitted using a linear combination of the following variables:

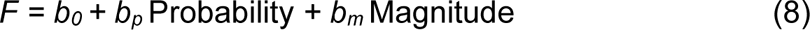

where Probability and Magnitude are the probability and magnitude of the rewards, respectively, as indicated by the pie chart. *b*_0_ is the intercept. If *b_p_* and *b_m_* were not zero at *P* < 0.05, the discharge rates were regarded as being significantly modulated by that variable. These results have been previously reported (*23*).

### Population dynamics using principal component analysis

We analyzed neural activity during an identical 0.6 s duration from the sample onset (Exp. 1), item onset (Exp. 2), CUE onset (Exp. 3), and cue onset (Exp. 4). To obtain a time series of neural firing rates within this time period, we estimated the firing rates of each neuron for every 0.05 s time bin (without overlap) during the analysis periods. A Gaussian kernel was not used.

#### Regression subspace

We used a general linear model to determine how items and locations (Exp. 1), items and scenes (Exp. 2), delay and magnitude of rewards (Exp. 3), and the probability and magnitude of the rewards (Exp. 4) affect the activity of each neuron in the neural populations. Each neural population was composed of all the recorded neurons in each brain region.

#### Exp. 1

First, we set six visual items and four locations as categorical variables. We then described the average firing rate of neuron *i* at time *t* as a linear combination of the item and the location in each neural population:

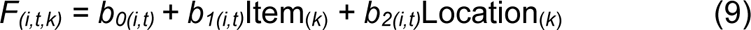

where *F_(i,t,k)_* is the average firing rate of neuron *i* at time t in trial *k,* Item_(*k*)_ is the types of items cued to the monkey in trial *k,* and Location_(*k*)_ *is* the types of locations cued to the monkey in trial *k*. The regression coefficients *b_0(i,t)_*, *b_1(i,t)_*, and *b_2(i,t)_* describe the degree to which the firing rates of neuron *i* depend on the mean firing rates (hence, firing rates independent of task variables, item, and location), the degree of firing rate in each item relative to the mean firing rates, and the degree of firing in each location relative to the mean firing rates, respectively, at a given time *t* during the trials. The interaction term is not included in the model.

In the analysis, we performed preference ordering for item and location in each neuron. Item_(*k*)_ and Location_(*k*)_ were rank-ordered items and locations, respectively, cued to the monkey in trial *k*. Items 1–6 and locations 1–4 were rank-ordered from the most preferred to least preferred, respectively, defined as the mean firing rate during the entire analysis time window from 0.08 to 0.6 s. This preference ordering did not change over time *t* for each neuron *n*.

#### Exp. 2

We first set eight items and two scenes as the categorical variables. We then described the average firing rate of neuron *i* at time *t* as a linear combination of the item and scene in each neural population:

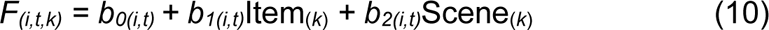

where *F_(i,t,k)_* is the average firing rate of neuron *i* at time t in trial *k,* Item_(*k*)_ is the types of items cued to the monkey in trial *k,* and Scene_(*k*)_ is the types of scene stimuli cued to the monkey in trial *k*. The regression coefficients *b_0(i,t)_*, *b_1(i,t)_*and *b_2(i,t)_* describe the degree to which the firing rates of neuron *i* depend on the mean firing rates (hence, firing rates independent of task variables, item and scene), the degree of firing rate in each item relative to the mean firing rates, and the degree of firing in each scene relative to the mean firing rates, respectively, at a given time *t* during the trials. The interaction term was not included in the model.

In the analysis, Item_(*k*)_ and Scene_(*k*)_ were the rank-ordered item and scene, respectively, cued to the monkey in trial *k*. Items 1 to 8 and Scenes 1 and 2 were rank-ordered from the most preferred to least preferred, respectively, defined as the mean firing rate during the whole analysis 0.6 s window after the item onset. This preference ordering did not change over time *t* for each neuron *n*.

#### Exp. 3

We first set the delay and magnitude as 0, 3.3, and 6.9 s and one and three drops of rewards, respectively, during the behavioral task. In the analysis, we normalized these values from 0 to 1 divided by the maximum values in each: 0, 0.48, and 1 for delay, and 0.33, 0.66, and 1 for magnitude. This is because these values affect the extent of the regression subspace between two continuous variables. We then described the average firing rate of neuron *i* at time *t* as a linear combination of the delay and magnitude in each neural population:

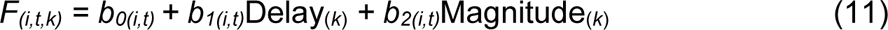

where *F_(i,t,k)_* is the average firing rate of neuron *i* at time t in trial *k,* Delay_(*k*)_ is the normalized delay to obtain a reward cued to the monkey in trial *k,* and Magnitude _(*k*)_ *is* the normalized number of reward drops cued to the monkey in trial *k*. The regression coefficients *b_0(i,t)_* to *b_2(i,t)_* describe the degree to which the firing rates of neuron *i* depend on the mean firing rates (hence, firing rates independent of task variables), delay in rewards, and magnitude of rewards, respectively, at a given time *t* during the trials.

#### Exp. 4

We first set the probability and magnitude as 0.1 to 1.0 and 0.1 to 1.0 mL, respectively. We did not normalize these values because they were originally prepared from 0 to 1 originally. We then describe the average firing rate of neuron *i* at time *t* as a linear combination of probability and magnitude in each neural population:

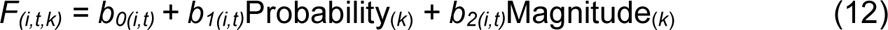

where *F_(i,t,k)_* is the average firing rate of neuron *i* at time t in trial *k,* Probability_(*k*)_ is the probability of the reward cued to the monkey in trial *k,* and Magnitude_(*k*)_ *is* the magnitude of the reward cued to the monkey in trial *k*. The regression coefficients *b_0(i,t)_* to *b_2(i,t)_* describe the degree to which the firing rates of neuron *i* depend on the mean firing rates (i.e., firing rates independent of task variables), probability of rewards, and magnitude of rewards, respectively, at a given time *t* during the trials.

We used the regression coefficients (i.e., the regression table in the case of ANOVA) described in Eq. 9–12 to identify how the dimensions of the neural population signals were composed of information related to the item and location (Exp. 1), item and scene (Exp. 2), delay and magnitude (Exp. 3), and probability and magnitude (Exp. 4) as aggregated properties of individual neural activity. In this step, an encoding model is constructed in which the regression coefficients are explained by a temporal structure in the neural modulation of two categorical variables (Exps. 1 and 2), or two continuous variables (Exps. 3 and 4) at the population level. Our procedures involve targeted dimensionality reduction using the regression subspace and are aimed at describing neural modulation dynamics (*29*).

#### Principal component analysis

We used PCA to identify the dimensions of the neural population signal in orthogonal spaces composed of two variables in each neural population of the four experiments. For each neural population, we first prepared a two-dimensional data matrix *X* of size *N_(n)_*×*M _(C_*_×*T)*_. The regression coefficient vectors *b_1(i,t)_* and *b_2(i,t)_* in Eq. 9–12, whose rows correspond to the total number of neurons (*n*) in each neural population and columns correspond to *C,* the total number of conditions (that is, 10: six items and four locations in Exp. 1, 10: eight items and two scenes in Exp. 2, 2: delay and magnitude in Exp. 3, and 2: probability and magnitude in Exp. 4), and T is the total number of analysis windows (i.e., 0.6 s divided by the window size bin, 0.05 s, 12 bin). A series of eigenvectors was obtained by applying PCA once to the data matrix *X* in each neural population. The PCs of this data matrix are vectors *v_(a)_* of length *N_(n)_* and the total number of recorded neurons if *M _(C_*_×*T)*_ > *N_(n)_*; otherwise, the length is *M _(C_*_×*T)*_. PCs were indexed from the principal components and explained the most to least variance. The eigenvectors were obtained using the prcomp () function in R software. We did not include the intercept term *b_0(i,t)_* to focus on the neural modulation by the variables of interest.

#### Eigenvectors

When we applied PCA to data matrix *X*, we decomposed the matrix into eigenvectors and eigenvalues. Each eigenvector had a corresponding eigenvalue. In our analysis, the eigenvectors at time *t* represented a vector, for example, in the space of delay and magnitude in Exp. 3. The eigenvalues at time *t* for the delay and magnitude were scalars, indicating the extent of variance in the data in that vector. Thus, the first PC was the eigenvector with the highest eigenvalue. We analyzed the eigenvectors for the top two PCs (PC1 and PC2) in the following analyses to describe the geometry in the most predominant dimension. PCA was applied once to each neural population; thus, the total variance contained in the data differed among the neural populations.

#### Shuffle control for PCA

To examine the significance of the population structures described by PCA, we performed three shuffle controls. The two-dimensional data matrix *X* was randomized by shuffling in three ways. In shuffled control 1, matrix *X* was shuffled by permutating the allocation of neuron *n* at time *i*. This shuffle provided a data matrix *X* of size *N_(n)_*×*M _(C_*_×*T)*_, eliminating the temporal structure of neural modulation by condition *C* in each neuron but retaining the neural modulations at time *t* at the population level. In shuffled control 2, matrix *X* was shuffled by permutating the allocation of time *i* in each neuron *n*. This shuffle provided a data matrix *X* of size *N_(neuron)_*×*M _(C_*_×*T)*_, eliminating the neural modulation structure under condition *C* maintained in each neuron but retaining the neural modulation in each neuron at the population level. In shuffled control 3, matrix *X* was shuffled by permutating the allocation of both time *i* and neuron *n*. In these three shuffle controls, matrix *X* was estimated to be 1,000 times. PCA performance was evaluated by constructing the distributions of the explained variances for PC1 to PC12. The statistical significance of the variances explained by PC1 and PC2 was estimated based on the 95th percentile of the reconstructed distributions of the explained variance or bootstrap standard errors (i.e., standard deviation of the reconstructed distribution). We note that because the significant dimensions of neural populations dynamics differed the 10 neural populations, we analyzed the neural dynamics at the top two dimension, PC1 and 2.

#### Analysis of eigenvectors

We evaluated the characteristics of the eigenvectors for PC1 and PC2 in each neural population in terms of vector angle, size, and deviation. The eigenvectors were evaluated for each of the task parameters described above: item and location in Exp. 1, item and scene in Exp. 2, delay and magnitude in Exp. 3, and probability and magnitude in Exp. 4. The angle is the vector angle from the horizontal axis from 0° to 360° against the main PCs. The size is the length of the eigenvector. The deviation is the difference between the vectors. The deviation from the mean vector for each neural population was estimated. These three eigenvector characteristics were compared among the populations at *P* < 0.05, using the Kruskal–Wallis test and Wilcoxon rank-sum test with Bonferroni correction for multiple comparisons. The vector during the first 0.1 s was extracted from these basic analyses.

To evaluate the neural population geometry using their selected feature, we estimated the accumulated angle difference weighted by the deviance:

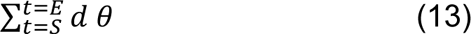

where the *d* is deviation between the vectors at times t and t+1, *θ* is the angle difference between vectors at times t and t+1, *S* is zero, and E is the time period to stop the estimation, i.e., 0.6 s. This index is analogous to the rotational force accumulated over time. If the value of the accumulated angle difference was close to zero, the population geometry was stable, such as a straight or non-dynamic structure, that is, it remained at some point in the PC1-2 plane.

To quantitatively evaluate the trajectory geometry, we used the Lissajous curve function, which describes any geometric pattern in a plane using *F(x,y)*:

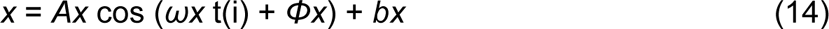

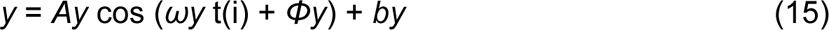

where ω and Φ represent cycle of the rotation and their deviance as a function of time, t(i). *Ax* and *Ay* represent the amplitudes of the trajectory, whereas *bx* and *by* represent the location of the trajectory. For *ω*, 3.33 π indicates that one cycle since the analysis window is 0.6 s. *Φ* is 0 to 2 π for one cycle. We estimated *ωx, Φx*, *bx, ωy,* and *Φy, by* parameters by estimating maximum loglikelihood of the model. Nonlinear least squares in the nls() function in the R program was used. A time series of eigenvectors for PC1 and PC2 in a 0.05 s analysis windows (12 data points) were used with a sliding average between three time points (hence, 0.15 s time resolution).

#### Bootstrap resampling and clustering using feature-based parameters

We estimated Σ*d θ*, mean *d*, rotational speed Σθ/0.1s, and *d*_s-e_, such as start to end distance using a parametric bootstrap resampling method (*65*). In each neural population, the neurons were randomly resampled in duplicate, and a data matrix *X* of size *N_(neuron)_*×*M _(C_*_×*T)*_ was obtained. PCA was applied to the data matrix *X.* The time series of eigenvectors was obtained, and these four features were estimated from the neural trajectory. This resampling was conducted 1,000 times in each neural population, and the distributions of these four parameters were obtained.

Following bootstrap resampling, we applied clustering of these parameters based on PCA and a dendrogram across the replicates in the 10 brain regions, such as 20,000 replicates (10 brain regions times two conditions times 1,000 replicates). Based on this clustering, proportion of the identified clusters in each brain region was estimated.

#### Bootstrap resampling and clustering based on Lissajous curve parameters

The Lissajous curve parameters for the replicated trajectory were estimated using a bootstrap resampling method (*65*). In each neural population, the neurons were randomly resampled in duplicate, and a data matrix *X* of size *N_(neuron)_*×*M _(C_*_×*T)*_ was obtained. PCA was applied to the data matrix *X.* The time series of eigenvectors was obtained for PC1 and PC2, which describe the trajectory, and the fitted parameters using the Lissajous curve function were estimated using the nls() function in R program. This resampling was conducted 1,000 times in each neural population, and the distributions of the Lissajous parameters were obtained.

Following bootstrap resampling, we applied clustering of these parameters based on PCA and a dendrogram across the replicates in the 10 brain regions, such as 20,000 replicates (10 brain regions times two conditions times 1,000 replicates). In this process, the omega ratio (*ωx*/*ωy*) and phi difference (*Φx*-*Φy*) were also used, in addition to the *ωx*, *ωy*, *Φx,* and *Φy*. Based on this clustering, proportion of the identified clusters in each brain region was estimated. We used the median of the estimated parameter in a cluster to describe the trajectory geometries.

## Acknowledgments

The authors thank Takashi Kawai, Ryo Tajiri, Yoshiko Yabana, Yuki Suwa, and Shiho Nishino for their technical assistance. Monkeys FU was provided by NBRP "Japanese Monkeys" through the National Bio Resource Project of the MEXT, Japan. Funding: This research was supported by JSPS KAKENHI (Grant Numbers JP:15H05374, 22H04832), JST Moonshot R&D JPMJMS2294 (H.Y.), and the National Natural Science Foundation of China (Grant 32271088) (Y.N.).

## Author Contributions

H.Y. conceptualized the study. H.Y., Y.N., T.M., O.H., and J.K. designed the experiments. H.Y., H.C., J.K., Y.I., and T.M. performed the experiments. H.Y. and Y.T. developed the analytical tools. H.Y., H.C., J.K., and Y.H. analyzed the data. H.Y., H.C., and J.K. wrote the manuscript. All authors edited and approved the final version of the manuscript.

## Data availability

All data and analysis codes used in this study are available in the supporting files.

Exp.1: Analysis code: Analy_mainchen.r and AnalyCue_PCAchen.r

Data: ChenMatData_All1506_inter_ranked.csv for 50 ms time bin
Chen_CellListAll1506.csv for cell type list

Exp. 2: Analysis code, Analy_mainkuni.r and AnalyCue_PCAkuni.r

Data: KuniMatData_All116_nointer_ranked.csv for 50 ms time bin

Exp. 3: Analysis code, Analy_mainhori.r and AnalyCue_PCAhori.r

Data: HoriMatData_All150.csv for 50ms time bin

Exp. 4: Analysis code, Analy_mainyamadai.r and AnalyCue_PCAyamada.r

Data: MatDataPCAProMag_50ms2021Sep09.txt for 50 ms data
CellList_DSVSOFCmOFC2019_0116.csv for cell type list

Common code for all experiments (1 to 4):

ParaData_BS1k50ms3ave.csv: Bootstrap replicate Data for the selected geometric features.
ParaData_BS1k50ms3aveLissague3final.csv: bootstrap replicate Data for Lissajous curve.
Analysis code for PCA: Analy_pca1.0final.r.

---

**Fig. S1.**
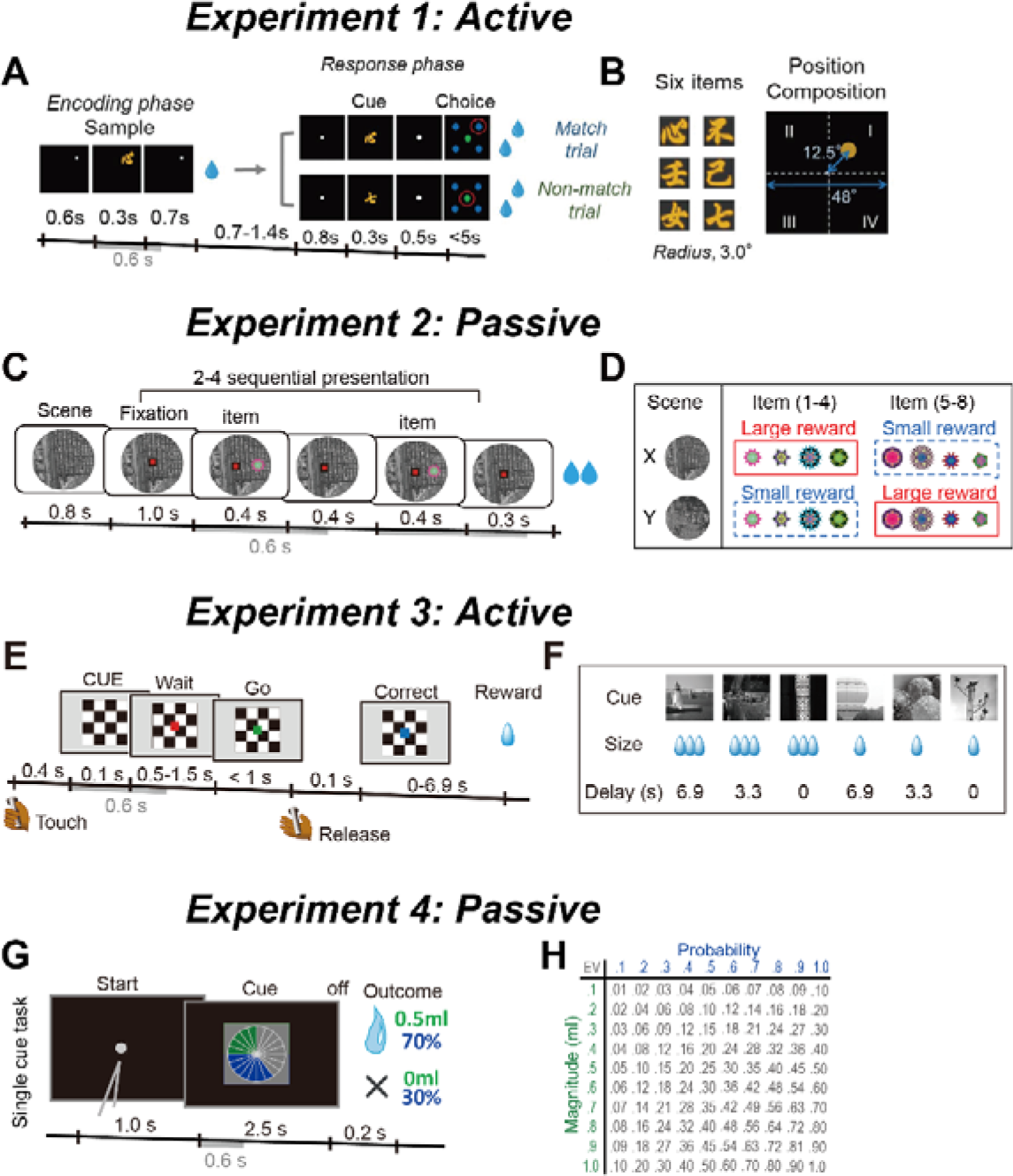
Behavioral tasks. **A,** Sequence of events during the ILR task in Exp. 1. The cue stimulus during the response phase was the same as the sample stimulus during the encoding phase of the match trial, whereas the two stimuli differed in the nonmatch trial. Neural activity was analyzed during 0.6 s after sample onset (gray bar). During the task, the monkeys actively responded or did not respond by making a choice and performing the following action. **B,** Six visual stimuli and spatial composition of the sample stimulus. **C,** Sequence of events during the passive viewing task in Exp. 2. During fixation, the visual items were presented sequentially within two to four times. These visual items were associated with reward or no-reward outcomes in other behavioral contexts during the learning trials. During the task, the monkeys were not required to respond, except for fixation to the center (passive). Neural activity was analyzed during 0.6 s after object onset (gray bars). **D,** Eight visual items were divided into two groups (items 1–4 and 5–8). In each scene (e.g., scene X), one group was associated with a large reward and the other with a small reward. This reward association was reversed in the other scene (e.g., scene Y). In each trial, two items appeared as shown in **C**, one in items 1–4 and the other in items 5–8 as a random combination. **E,** Sequence of events during the delayed reward task in Exp. 3. During the task, the monkeys actively responded to the GO signal by releasing the lever (active). Neural activity was analyzed during 0.6 s after cue onset (gray bar). **F,** During the task, six visual items indicated the forthcoming reward size and delay duration to the reward after the bar release. **G,** Sequence of events during the single-cue task in Exp. 4. A single visual pie chart with green and blue segments was presented to the monkeys. During the task, the monkeys were not required to respond, except for fixation to the center during the start (passive). Neural activity was analyzed during 0.6 s after cue onset. **H,** Payoff matrix: Each of the magnitudes was fully crossed with each of the probabilities, resulting in a pool of 100 lotteries.

**Fig. S2.**
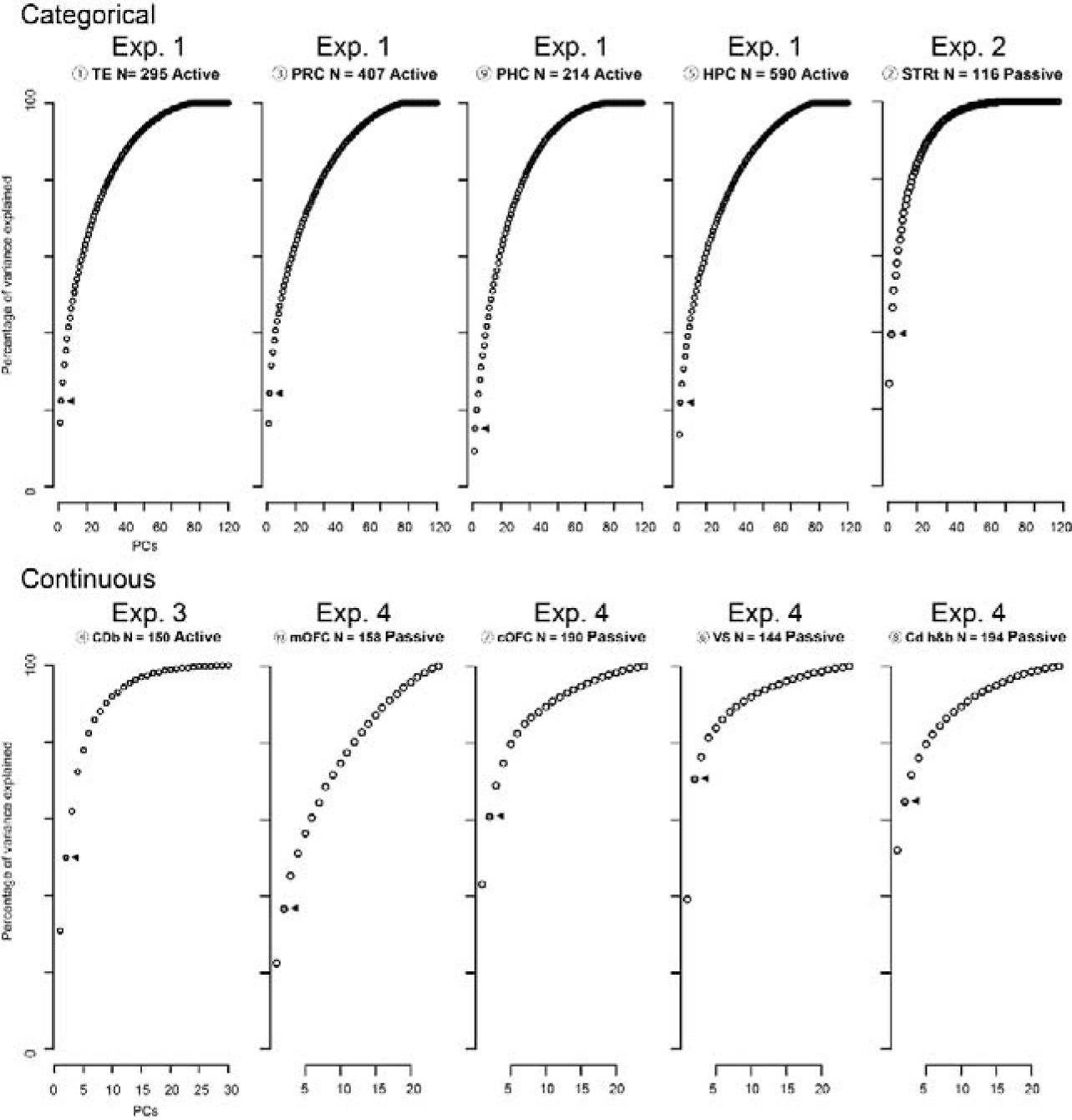
Variances explained by PCA in each neural population. Cumulative variance explained by PCA in each neural population in Exp. 1 to 4. The 0.05 s time bin was used for the analysis. In Exp. 1 and Exp. 2, categorical task parameters were used. In Exp. 3 and Exp. 4, continuous task parameters were used. Triangle indicates the variance explained by the first two PCs.

**Fig. S3.**
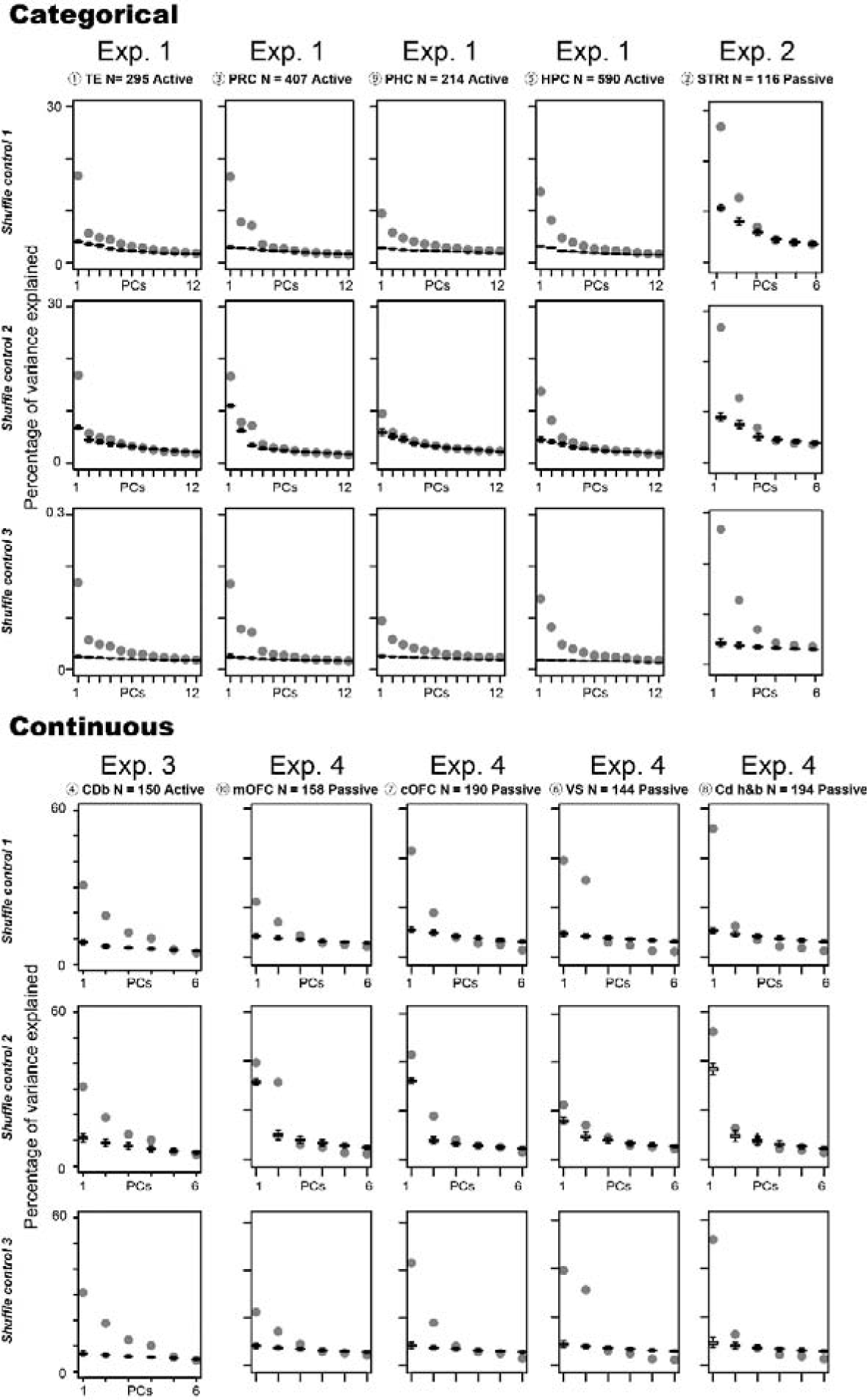
Variances explained by PCA under shuffle control. Boxplot of variances explained by PCA under the three shuffled conditions (see Methods section for details). The plot was not cumulative. A boxplot was made with 1,000 repeats of the shuffle for each condition. Gray plots indicate the percentage of variance explained by PCA. Results using 0.05 s bin data are shown.

**Fig. S4.**
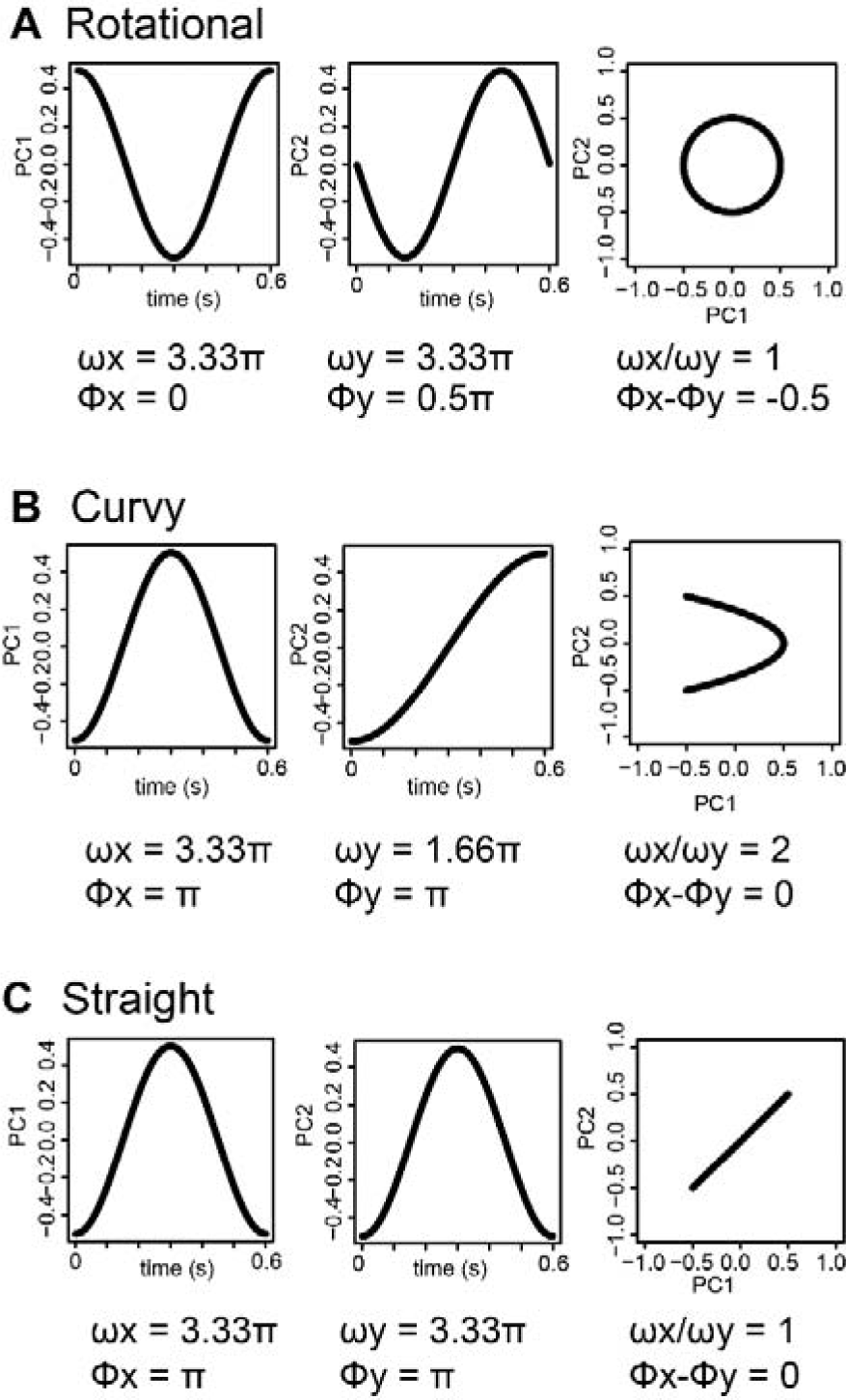
Examples of Lissajous curve represented with the parameter combination. Rotational, curvy, and straight dynamics are shown against *ω* ratio and *Φ* difference. In this study one cycle is defined with 3.33 π for *ω* during 0.6 s analysis period (3.33 π × 0.6 = 2 π in the Lissajous function, *A* cos(*ω* t(i) + *Φ*) + *b*). *Φ* from 0 to 2 π defines the phase. *A* and *b* denote the size and location of the curve, respectively. Combination of *ω* ratio and *Φ* difference between x and y determines the shape of trajectory. For example, straight geometry is defined as the same *ω* and the same *Φ*. Rotational dynamics is defined as the same *ω* and not identical *Φ*. Curvy dynamics is defined as the different *ω* and the same *Φ*.

**Supplementary Table S1.**
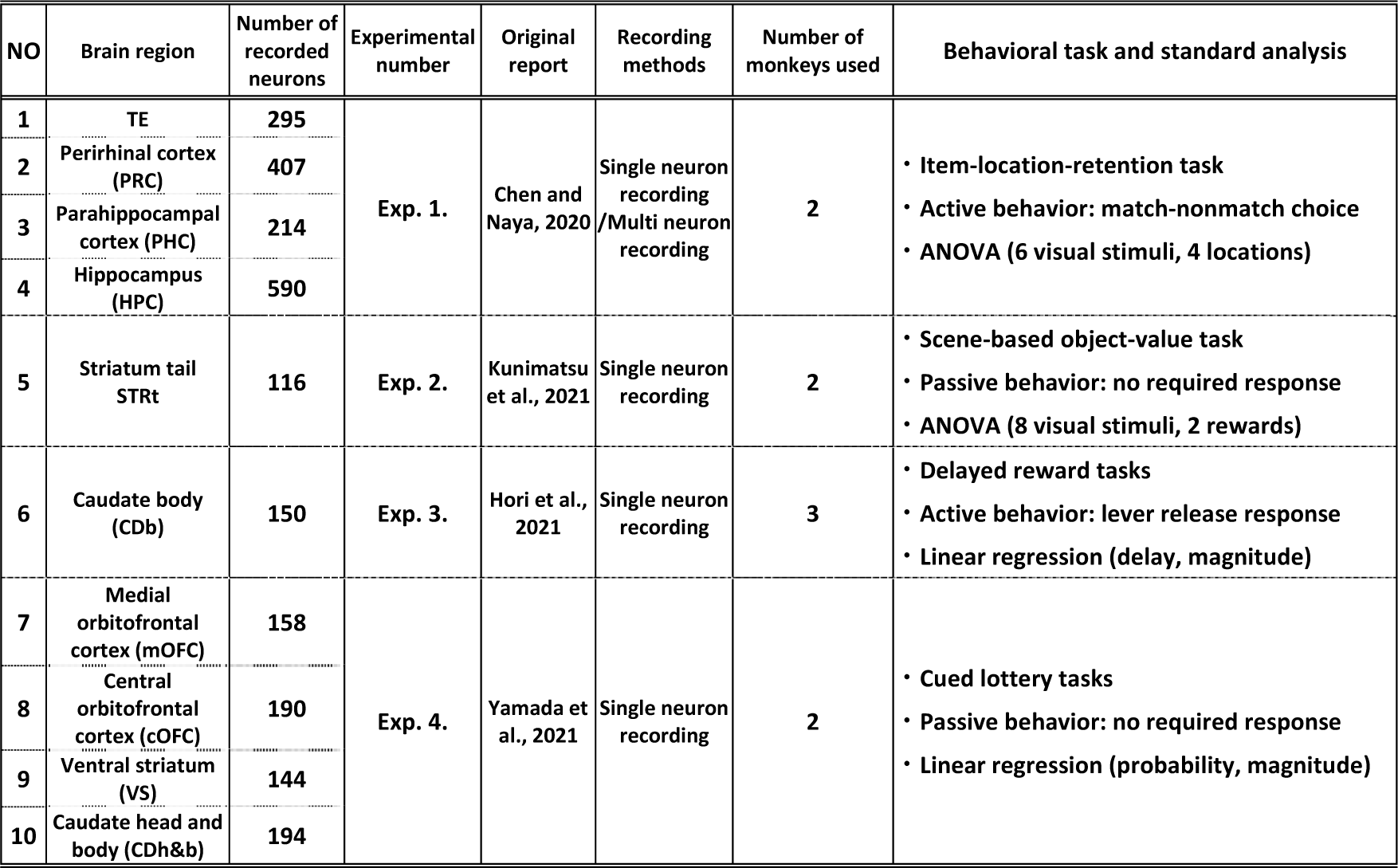
Summary of the data set and standard analysis methods. For the standard analysis, we analyzed visual response within a short time (0.6 s). Neural modulations were detected at the same significance level (P < 0.05) using a general linear model (ANOVA in Exps. 1 and 2 and the linear regression in Exp. 3 and 4). See each reference for the results based on the standard analysis using general linear model.

| NO | Brain region | Number of recorded neurons | Experimental number | Original report | Recording methods | Number of monkeys used | Behavioral task and standard analysis |
| --- | --- | --- | --- | --- | --- | --- | --- |
| 1 | TE | 295 | Exp. 1. | Chen and Naya, 2020 | Single neuron recording / Multi neuron recording | 2 | • Item-location-retention task<br>• Active behavior: match-nonmatch choice<br>• ANOVA (6 visual stimuli, 4 locations) |
| 2 | Perirhinal cortex (PRC) | 407 |  |  |  |  |  |
| 3 | Parahippocampal cortex (PHC) | 214 |  |  |  |  |  |
| 4 | Hippocampus (HPC) | 590 |  |  |  |  |  |
| 5 | Striatum tail STRt | 116 | Exp. 2. | Kunimatsu et al., 2021 | Single neuron recording | 2 | • Scene-based object-value task<br>• Passive behavior: no required response<br>• ANOVA (8 visual stimuli, 2 rewards) |
| 6 | Caudate body (CDb) | 150 | Exp. 3. | Hori et al., 2021 | Single neuron recording | 3 | • Delayed reward tasks<br>• Active behavior: lever release response<br>• Linear regression (delay, magnitude) |
| 7 | Medial orbitofrontal cortex (mOFC) | 158 | Exp. 4. | Yamada et al., 2021 | Single neuron recording | 2 | • Cued lottery tasks<br>• Passive behavior: no required response<br>• Linear regression (probability, magnitude) |
| 8 | Central orbitofrontal cortex (cOFC) | 190 |  |  |  |  |  |
| 9 | Ventral striatum (VS) | 144 |  |  |  |  |  |
| 10 | Caudate head and body (CDh&b) | 194 |  |  |  |  |  |

